# Lipid droplet lipolysis in POMC neurons regulates energy homeostasis in a sex-specific manner

**DOI:** 10.64898/2026.09.16.752127

**Authors:** Danie Majeur, Romane Manceau, Frédérick Boisjoly, Lewis Rhys Depaauw-Holt, Marie-Flore Fourn, Josephine Louise Robb, Demetra Rodaros, Khalil Bouyakdan, Stephanie Fulton, Ciaran Murphy-Royal, Thierry Alquier

## Abstract

The hypothalamus is a central regulator of glucose and energy homeostasis, with arcuate nucleus (ARC) neurons orchestrating these processes. Agouti-related peptide (AgRP) and pro-opiomelanocortin (POMC) neurons integrate metabolic cues to control feeding behavior and systemic metabolism. Among these cues, fatty acids (FA) have emerged as key modulators of ARC neuronal activity. While neuronal sensing of circulating FA has begun to be defined, the contribution of FA derived from endogenous lipid stores to ARC neuron function and energy homeostasis remains largely unexplored. We recently identified lipid droplets (LD) as regulated FA reservoirs that control FA availability and metabolism in orexigenic AgRP neurons, thereby modulating their activity and regulating feeding. This prompted us to investigate whether LD-derived FA similarly regulate POMC neuron function. To this end, we targeted adipose triglyceride lipase (ATGL), which catalyzes the first committed step of LD lipolysis, in POMC neurons. We show that LD are present in POMC neurons under basal conditions both in vitro and in vivo, and that pharmacological or genetic inhibition of ATGL increases LD abundance. ATGL deficiency enhances spontaneous firing of POMC neurons and leads to reduced body weight, fat and lean mass in males, but not females. Consistent with enhanced glucose metabolism, ATGL loss lowers glycemia and insulinemia and increases carbohydrate utilization in chow-fed males. In contrast, ATGL deficiency does not alter metabolic adaptations to cold exposure, fasting or diet-induced obesity in either sex. Collectively, these findings establish ATGL-dependent LD lipolysis in POMC neurons as a previously unrecognized, sex-dependent regulator of energy homeostasis.

## INTRODUCTION

The hypothalamus plays a central role in maintaining energy homeostasis by integrating hormonal and nutrient-derived signals to coordinate feeding behavior and metabolism. Among these signals, fatty acids (FA) have emerged as important metabolic cues that regulate hypothalamic neuronal activity and whole-body energy balance. Numerous studies have demonstrated that circulating or exogenous FA modulate the activity of hypothalamic neurons, including agouti-related peptide (AgRP) and pro-opiomelanocortin (POMC) neurons of the arcuate nucleus (ARC), thereby influencing feeding behavior and energy homeostasis^1^. These effects depend on both the type of FA (short vs. long chain) and neuronal populations^2–4^. Current evidence indicates that extracellular FA are detected through complementary mechanisms involving the FA receptor FFAR1^2,5,6^, FA uptake and transport via lipoprotein lipase and CD36 ^7,8^ and intracellular metabolic processing^3,9^.

Beyond the sensing of circulating FA, increasing evidence suggests that endogenous intracellular FA are also critical regulators of hypothalamic neuron function. These FA may originate from *de novo* lipogenesis or from intracellular lipid stores, and their metabolic partitioning between esterification and mitochondrial FA oxidation (FAOx) influences neuronal excitability and systemic metabolism. In AgRP neurons, disruption of FA esterification into glycerolipids^10^, mitochondrial dynamics and carnitine palmitoyltransferase 1 (CPT1)-dependent FAOx^11,12^ impair neuronal activity, glucose sensing and feeding behavior. Likewise, partial disruption of oxidative phosphorylation in POMC neurons increases FAOx and reduces neuronal activity^13^, whereas deletion of fatty acid synthase, responsible for *de novo* FA synthesis, enhances POMC neuron activity and promotes a lean phenotype with improved glucose homeostasis^14^. Together, these studies establish intracellular FA metabolism as a key regulator of hypothalamic neuronal function. However, they have primarily focused on FA synthesis and mitochondrial oxidation, leaving unresolved whether FA mobilized from intracellular lipid stores similarly regulate POMC neuron activity and energy homeostasis.

Lipid droplets (LD) are dynamic organelles composed primarily of triglycerides and cholesteryl esters that buffer intracellular FA availability by storing excess lipids and releasing them through regulated lipolysis^15–17^. Rather than serving as inert lipid depots, LDs are increasingly recognized as dynamic hubs that coordinate lipid and energy metabolism, organelle communication, stress responses and cell fate^16,17^. The rate-limiting step of LD lipolysis is catalyzed by adipose triglyceride lipase (ATGL), which releases FA that can subsequently be directed toward oxidation, membrane lipid synthesis or signaling pathways^18^. Although historically overlooked in neurons, recent studies reported that triglycerides and LD are present in neurons *in vitro*^19,20^ and LD lipolysis influences neuronal metabolism and activity^21–24^. Thus, LD lipolysis represents a potential mechanism linking intracellular LD to neuronal metabolism and function.

Consistent with this concept, we recently demonstrated that LD are present in AgRP/Neuropeptide Y (NPY) neurons and that pharmacological inhibition or genetic deletion of ATGL promotes LD accumulation in these cells^25^. Moreover, deletion of ATGL restricted to ARC neurons altered energy expenditure, body temperature and feeding responses during cold exposure^25^. Notably, these metabolic alterations were only partially reproduced in mice with ATGL deletion specifically in AgRP neurons, suggesting that additional ARC neuronal populations contribute to the physiological actions of neuronal LD lipolysis. Given the established role of intracellular FA metabolism in POMC neurons, we hypothesized that ATGL-mediated mobilization of FA from LD regulates POMC neuron activity and energy homeostasis. To test this hypothesis, we generated mice lacking ATGL specifically in POMC neurons (POMC^ATGL^KO) and investigated the contribution of LD lipolysis to neuronal activity and whole-body glucose and energy metabolism. Loss of ATGL increased POMC neuron firing which was associated with lower body weight, lean and fat mass, and enhanced carbohydrate utilization in chow-fed males but not females. POMC^ATGL^KO males, but not females, had reduced food intake after a fast, suggesting increased satiety. Metabolic responses during fasting, cold exposure and diet-induced obesity were not affected by ATGL deficiency. Our findings identify ATGL-dependent LD lipolysis as a previously unrecognized, sex-specific regulator of POMC neuron excitability and metabolic homeostasis.

## RESULTS

### Lipid droplet lipolysis is regulated by ATGL in POMC neurons

We previously showed that LD are present in AgRP/NPY neurons in which ATGL-mediated LD lipolysis is required for their activity and feeding responses^25^. Thus, we investigated whether LD are present and regulated by ATGL in POMC neurons. We first assessed if LD are present in primary POMC neurons derived from POMC-GFP mice under basal conditions. Our results show that, under basal conditions in vitro, approximately 60% of primary POMC neurons accumulate LD, with an average of ∼1 LD per neuron (Fig. 1a-d). ATGL inhibition with ATGListatin significantly increased the percentage of LD^+^ POMC neurons and the number of LD/neuron (Fig.1a-d). Next, we generated an ATGL loss-of-function mouse model using POMC-Cre and ATGL floxed mice crossed with the POMC-GFP strain to visualize POMC neurons (Fig. S1a). Using qPCR and RNAscope, we validated that ATGL expression is reduced in POMC^ATGL^ KO mice compared to POMC^ATGL^ CRE control littermates (Fig. S1b-e). The number of GFP^+^ POMC neurons was not affected by loss of ATGL (Fig. S1f-h) suggesting cell survival is not influenced.

**Figure 1.**
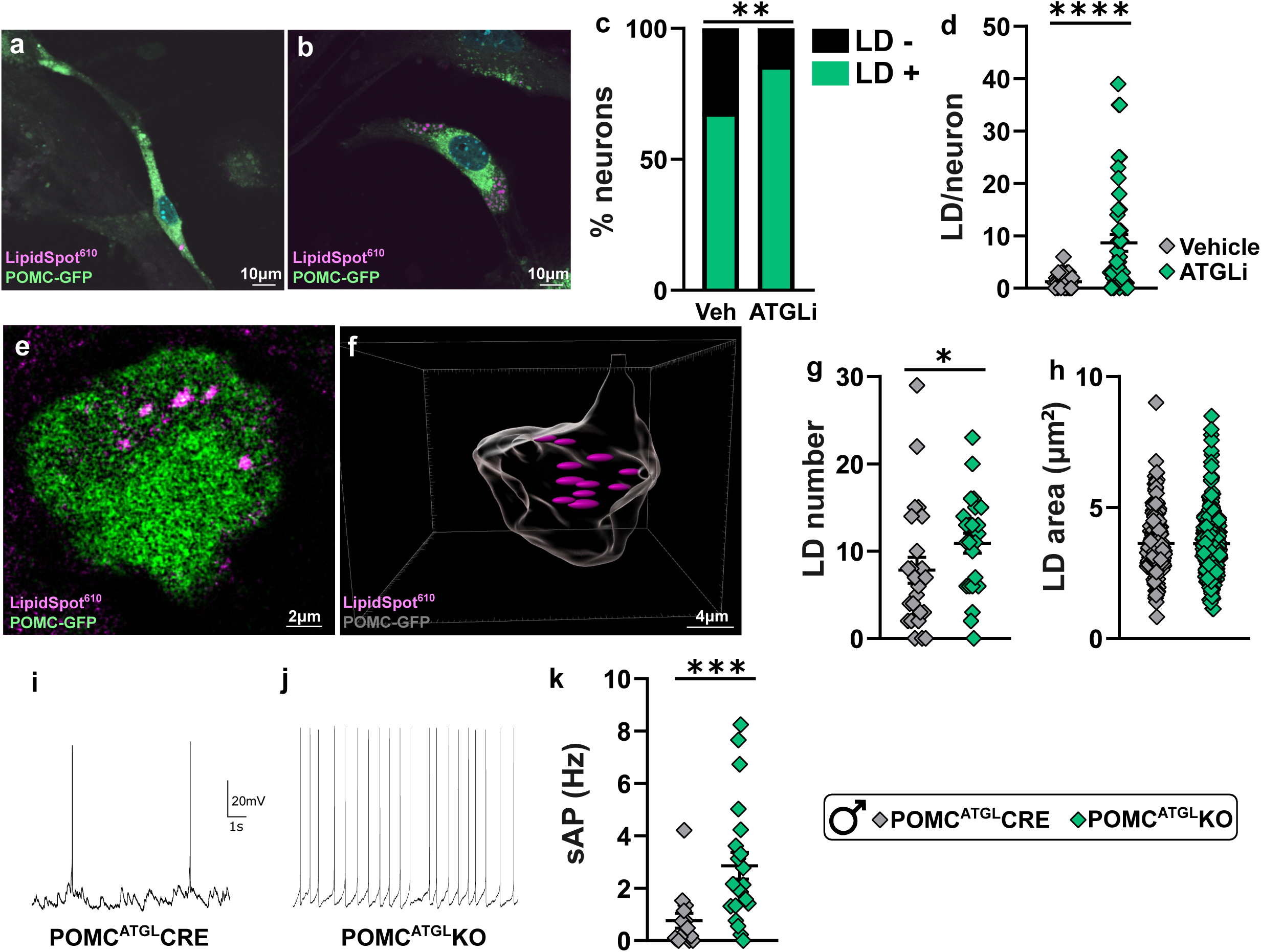
ATGL regulates lipid droplets in POMC neurons and neuronal activity. LD imaging with LipiSpot^610^ in primary POMC neurons (derived from POMC-GFP pups) treated with vehicle (DMSO) (**a**) or the ATGL inhibitor, Atglistatin (ATGLi, 50 µM) (**b**). Proportion of LD-positive (LD^+^) and negative (LD^-^) POMC-GFP neurons (**c**) and number of LD in POMC-GFP positive neurons (**d**) with vehicle (DMSO) or ATGLi for 24 h. N = 37-43 neurons/conditions. Representative 2D confocal image of LipidSpot^610^ staining in a POMC-GFP neuron (z-steps of 0.3 µm, N = 24-25 neurons from 2 POMC^ATGL^KO and 2 POMC^ATGL^CRE male mice) (**e**) and corresponding 3D reconstruction (**f**). POMC LD number (**g**) and LD area (**h**) in brain sections of POMC^ATGL^CRE and POMC^ATGL^KO male mice. Sample traces of spontaneous action potentials (sAP) in POMC^ATGL^CRE (**i**) and POMC^ATGL^KO (**j**) and corresponding quantification of frequency of sAP (**k**) in male mice. N = 15-21 neurons from 3 POMC^ATGL^KO and 3 POMC^ATGL^CRE mice. ATGLi; Atglistatin, Veh; Vehicle, LD; Lipid Droplet. Data are represented as mean ± SEM. Fisher’s exact test (**c**), Student’s t-test (**d, h, k**) or Mann-Whitney test (**g**). *p<0.05, **p<0.01, ***p<0.001,****p<0.0001.

LD number and area were measured in POMC^ATGL^CRE and KO male mice using Lipidspot^610^ and confocal imaging with 3D reconstruction of POMC neuron soma. POMC neurons had 10 LD on average in POMC^ATGL^CRE mice, with a significant increase in LD number in POMC^ATGL^KO mice (Fig. 1e-g). LD area was not changed in POMC^ATGL^KO mice compared to POMC^ATGL^CRE littermates (Fig. 1h). Taken together, these data show that POMC neurons accumulate LD in physiological conditions and that ATGL regulates LD content.

### Loss of ATGL in POMC neurons increases POMC neuron activity in males

Our previous study showed that loss of ATGL in AgRP neurons reduces their firing^25^. We thus investigated the impact of ATGL loss on POMC neuron activity using patch-clamp electrophysiology. Loss of ATGL in POMC neurons led to increased frequency of action potentials in POMC^ATGL^KO (2.86 ± 0.51 Hz) compared to POMC^ATGL^CRE (0.76 ± 0.28 Hz) male mice (Fig. 1i-k). The frequency of action potentials in POMC neurons from POMC^ATGL^KO females was not affected compared to control POMC^ATGL^CRE mice (Fig. S2a). These findings indicate that blockage of LD lipolysis increases POMC neuron activity in a sex-dependent manner, an effect opposite to the reduced activity of AgRP neurons we observed in AgRP^ATGL^KO mice^25^.

### ATGL deficiency in POMC neurons modulates energy homeostasis in a sex-specific manner

With evidence that ATGL regulates LD content and activity of POMC neurons in males, we next sought to investigate the effects of ATGL loss in POMC neurons on energy homeostasis in chow-fed mice. Our results show that loss of ATGL in POMC neurons leads to decreased body weight (Fig. 2a-b), lean and fat mass (Fig. 2c) with reduced subcutaneous white adipose tissue (Fig. 2d) in male mice. Importantly, lower body weight was not associated with reduced cumulative food intake over 8 weeks of standard chow diet (Fig. 2e). Moreover, these changes were not associated with alterations of femur length suggesting that overall growth was not affected in POMC^ATGL^KO males (Fig. S1i) or females (Fig. S2b). Based on the known role of POMC neurons on hypothalamus-pituitary axis activity^26,27^, we measured plasma corticosterone following acute restraint stress. POMC^ATGL^CRE and POMC^ATGL^KO male mice had similar corticosterone levels in basal conditions (19.04 ± 3.41 ng/mL POMC^ATGL^CRE vs. 17.42 ± 2.71 ng/mL POMC^ATGL^KO, N = 11-12, p>0.05) and after restraint stress (53.36 ± 5.64 ng/mL POMC^ATGL^CRE vs. 66.32 ± 5.90 ng/mL POMC^ATGL^KO, N = 10-12, p>0.05). As POMC neurons are key regulators of glucose homeostasis, we next assessed glucoregulatory responses in POMC^ATGL^CRE and KO males. Glycemia and insulinemia were reduced in POMC^ATGL^KO males (Fig. 2f,i). While glucose tolerance was not affected during a tolerance test, we found reduced glucose-induced insulin secretion in POMC^ATGL^KO males compared to control littermates (Fig. 2g-h, j-k). These findings thereby suggest increased insulin sensitivity in POMC^ATGL^ KO males. Interestingly, body weight and body composition were not affected in POMC^ATGL^ KO females compared to controls (Fig. S2c-e). This suggests that ATGL-dependent LD lipolysis in POMC neurons is required for proper control of whole-body energy and glucose homeostasis in male but not female mice.

**Figure 2.**
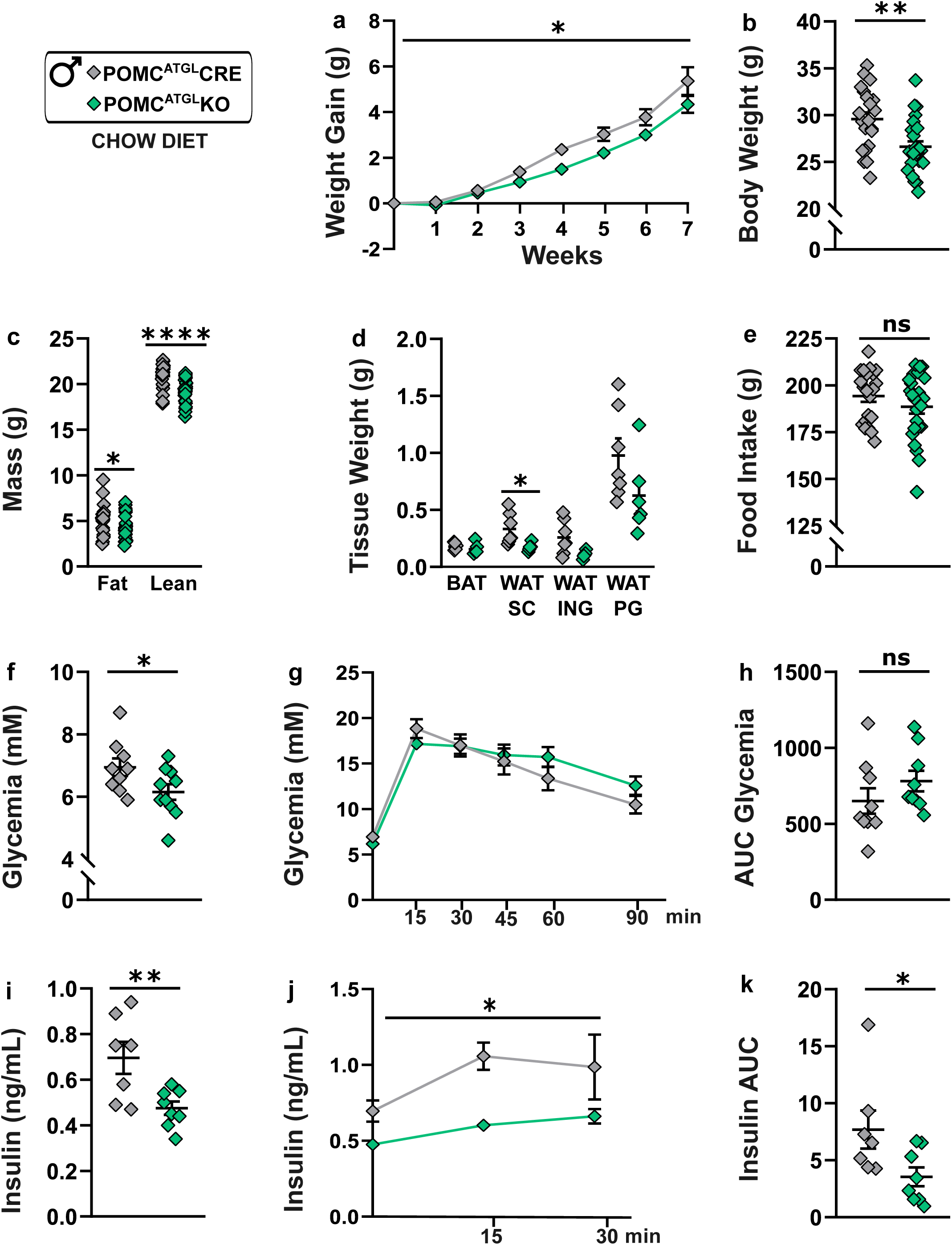
Loss of ATGL in POMC neurons affects body weight and glucose homeostasis. Body weight gain (**a**), body weight (**b**), body composition (**c**), fat depot weight (**d**) and cumulative food intake (**e**) in POMC^ATGL^KO and POMC^ATGL^CRE males after 8 weeks on standard chow diet. N = 21-25 mice/genotype (**a, b, c, e**) and N = 5-7 mice/genotype (**d**). Fasted (5 h) glycemia (**f**) and glycemia during an intra-peritoneal glucose tolerance test (**g-h**) in POMC^ATGL^KO and POMC^ATGL^CRE males. Fasted (5 h) insulin (**i**) and insulin during an intra-peritoneal glucose tolerance test (**j-k**) in POMC^ATGL^KO and POMC^ATGL^CRE males. N = 7-10 mice/genotype. AUC; Area under the curve. Data are represented as mean ± SEM. Student’s t-test (**b-f, h-i, k**) and two-way ANOVA with Sidak’s post-hoc test (**a, g, j**); * indicates a genotype effect while # indicates time effect. *p<0.05, **p<0.01, **** p<0.00001.

### Loss of ATGL in POMC neurons promotes carbohydrate utilization in chow-fed male mice

We next investigated if loss of ATGL in POMC neurons alters adaptive responses to metabolic challenges, particularly during fasting and cold exposure, conditions that are associated with increased ATGL expression in the ARC^25^. We first measured parameters of energy balance in metabolic cages in chow-fed conditions and during a fast-refeeding paradigm. In chow-fed conditions, energy expenditure (EE) (Fig. 3a,e) and locomotor activity (Fig. 3b,f) were not affected in POMC^ATGL^KO males in the light or dark period suggesting that the lower body weight in POMC^ATGL^KO males is not related to changes in EE or locomotor activity. POMC^ATGL^KO males showed an increase in respiratory quotient (RQ) (Fig. 3c,g) and a decrease in fatty acid oxidation (FAOx) (Fig. 3d,h) in fed conditions. During a fast, RQ, EE, FAOx and locomotor activity were not affected in POMC^ATGL^KO vs. POMC^ATGL^CRE males (Fig. 3a-h). Refeeding after the fast led to the expected rise of RQ and decrease of FAOx in POMC^ATGL^CRE and POMC^ATGL^KO males (Fig. 3c-d, g-h). However, POMC^ATGL^KO males showed an overall increase in RQ and decrease of FAOx compared to POMC^ATGL^CRE males (Fig. 3c-d, g-h). Meal pattern analysis showed that POMC^ATGL^KO males did not have the expected increase in total food intake following a fast as observed in POMC^ATGL^CRE mice (Fig. 3i), which was associated with no changes in meal number (Fig. 3j) but a decrease in meal size during the refeeding (Fig. 3k). With the exception of reduced FAOx during the fast, EE, RQ, locomotor activity, food intake and meal pattern were not affected in POMC^ATGL^KO females (Fig. S2f-p). Overall, these results suggest that loss of ATGL in POMC neurons shows a strong sex-specific effect, wherein ATGL loss in male mice promotes carbohydrate utilization in chow-fed conditions and reduces food intake following a fast.

**Figure 3.**
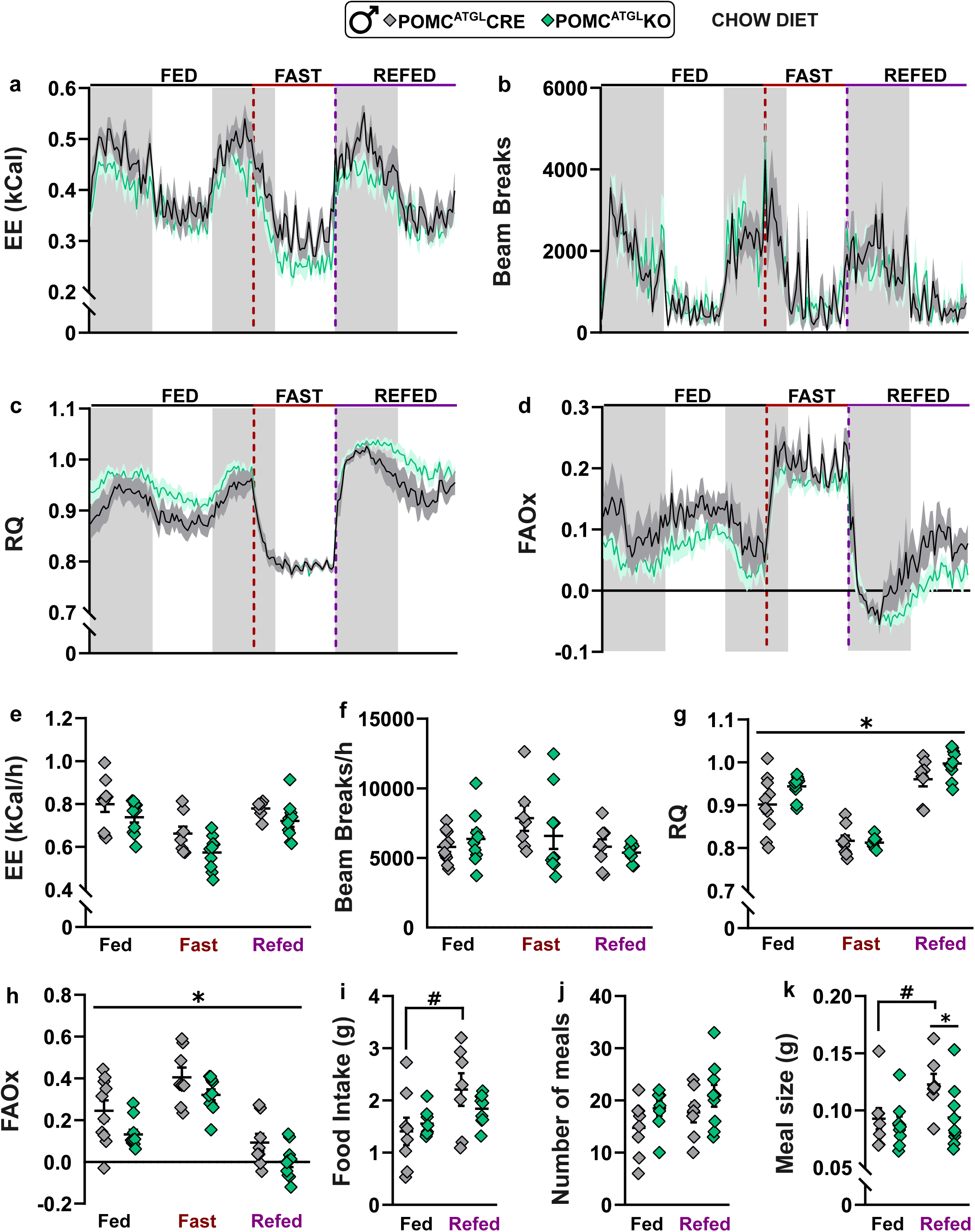
Parameters of energy balance in POMC^ATGL^KO males during a fast-refeeding paradigm. Metabolic parameters measured in metabolic cages including energy expenditure (EE) (**a, e**), locomotor activity (**b, f**), respiratory quotient (RQ) (**c, g**), and fatty acid oxidation (FAOx) (**d, h**) in POMC^ATGL^KO and POMC^ATGL^CRE male mice during 24 h in chow-fed conditions, a 16 h fast and 24 h refeeding. N = 10 mice/genotype. Food intake (**i**), number of meals (**j**) and meal size (**k**) in POMC^ATGL^KO and POMC^ATGL^CRE male mice during fed and refed conditions. N = 8-9 mice/genotype. Data are represented as mean ± SEM. Two-way ANOVA with Sidak’s post-hoc test (**a-k**); * indicates a genotype effect while # indicates time effect. * or ^#^ p<0.05.

Next, we assessed adaptive responses to cold exposure in POMC^ATGL^KO mice. POMC^ATGL^KO and CRE mice were exposed to cold (4°C) during 24 h while housed in metabolic cages. EE, locomotor activity, RQ, FAOx, core body temperature, food intake, meal number and meal size were similar during cold in POMC^ATGL^KO and CRE males (Fig. S3a-h) or females (Fig. S3i-p). Taken together, these data suggest that loss of ATGL in POMC neurons affects RQ and FAOx in chow-fed conditions with no changes in metabolic substrates utilization during fasting or cold exposure in males.

### Loss of ATGL in POMC neurons does not protect against diet-induced obesity

Based on our findings suggesting that loss of LD lipolysis in POMC neurons increases POMC neuron activity and favors carbohydrate utilization in male mice, we tested whether loss of ATGL protects from diet-induced obesity and associated glucose intolerance. To this end, POMC^ATGL^CRE and KO males were fed with a high-fat diet (HFD; 50 % and 33 % kcal derived from fat and sugar respectively) for 8 weeks. Under these diet conditions, body weight, body weight gain (Fig. 4a-b), food intake (Fig. 4c) and body composition (Fig. 4d-e) were similar in POMC^ATGL^KO and control littermates. Parameters of energy balance (EE, RQ, FAOx and activity), glycemia and tolerance to glucose were not altered in POMC^ATGL^KO mice during HFD (Fig. 4f-p). These results suggest that loss of ATGL from POMC neurons does not protect against diet-induced obesity. One possible explanation is that HFD feeding itself suppresses ATGL expression and/or activity in ARC neurons as described in adipose tissues^28^. In line with this idea, ATGL expression was reduced in the ARC of WT male mice fed with a HFD (Fig. S4a). In addition, ATGL expression in the ARC was not further decreased in POMC^ATGL^KO males compared to CRE controls fed with a HFD (Fig. S4b). Finally, POMC^ATGL^CRE and KO females fed with a HFD for 12 weeks showed similar changes in body weight (Fig. S4c), body composition (Fig. S4d-e), food intake (Fig. S4f), glycemia (Fig. S4g-h) or parameters of energy balance (Fig. S4i-l). This suggests that LD lipolysis in POMC neurons does not modulate responses to a HFD in males and females.

**Figure 4.**
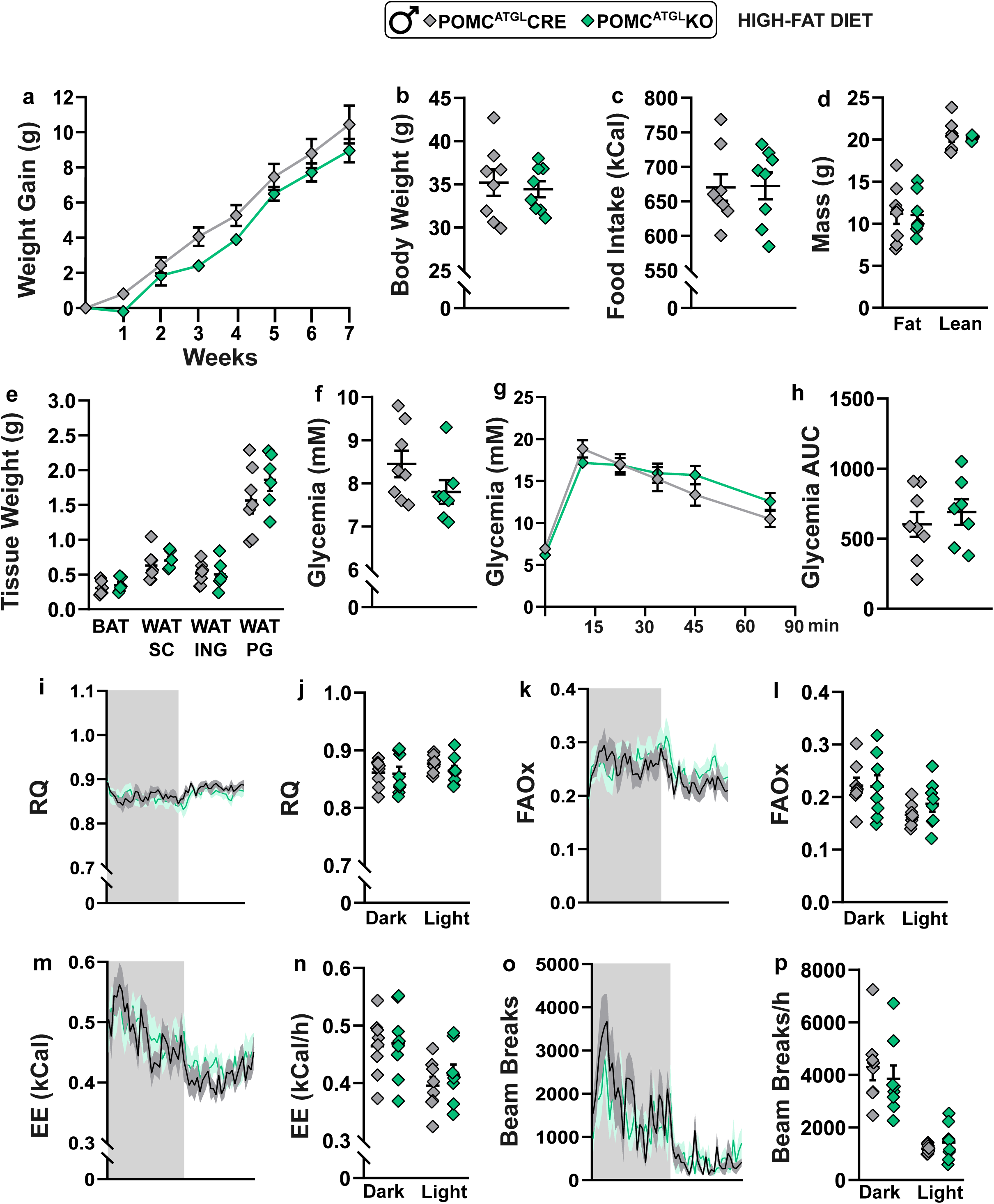
Diet-induced obesity is not affected by loss of ATGL in POMC neurons. Body weight gain (**a**), body weight (**b**), food intake (**c**), and body composition (**d**) of POMC^ATGL^KO and POMC^ATGL^CRE males after 8 weeks of HFD. Fat depot weight at sacrifice (**e**). Fasted (5h) glycemia (**f**) and glycemia (**g-h**) during an intra-peritoneal glucose tolerance test. Parameters of energy balance measured in metabolic cages including RQ (**i-j**), FAOx (**k-l**), EE (**m-n**), and locomotor activity (**o-p**) in POMC^ATGL^KO and POMC^ATGL^CRE male mice during 24 h. N = 8 mice/genotype. Data are represented as mean ± SEM. Two-way ANOVA with Sidak’s post hoc test (**a, g, i-p**) or Student’s t-test (**b-f, h**).

## DISCUSSION

Our data provides strong evidence that ATGL-dependent LD lipolysis in POMC neurons is required to maintain energy homeostasis in a sex-specific manner. We found that loss of ATGL in POMC neurons increases LD number *in vitro* and *in vivo*. Importantly, loss of ATGL increased POMC neuron activity in males but not females. Although loss of ATGL in POMC neurons did not affect energy homeostasis under states of negative energy balance (fasting and cold) or during HFD, it has notable sex-specific effects on energy homeostasis in normal chow-fed and refed conditions. We found that POMC^ATGL^KO males, but not females, had reduced body weight, lean mass and fat mass. This lean phenotype was associated with lower fasted blood glucose and insulin levels, reduced glucose-induced insulin secretion and increased RQ supporting increased insulin sensitivity and carbohydrate utilization. Additionally, food intake was blunted following a fast in POMC^ATGL^KO males. Together, these findings reveal an important sex-specific role for intracellular FA originating from LD lipolysis in POMC neurons in promoting neuronal activity and maintaining energy homeostasis.

Our findings indicate that LD number increases in primary POMC neurons *in vitro* and *vivo*. However, we did not detect any changes in LD size *in vivo* (Fig. 1g-h). This increase in LD number but not size with loss of ATGL in POMC neurons suggests that FA are preferentially stored in nascent LD rather than in preexisting LD. This phenotype may indicate a limited capacity of POMC neurons to increase LD size. Future studies are needed to determine whether differences in LD turnover or biogenesis contribute to increased LD number but not size in POMC neurons with loss of ATGL.

Our results show that POMC neurons in POMC^ATGL^KO males display increased frequency of spontaneous action potential (Fig. 1i) thereby suggesting that FA released by ATGL have an inhibitory effect on POMC neuron activity. While additional studies will be required to elucidate the underlying electrophysiological mechanisms, they may implicate alteration of ion channel activity by FA including voltage-gated Na⁺ channels or K_ATP_ channels^3,4^. Consistent with our data suggesting that LD-derived FA inhibit POMC neuron activity, loss of FA synthase also increases POMC neuron activity^14^. These findings thus suggest that *de novo* synthesis or release of FA from endogenous stores silences POMC neurons. We previously reported that ATGL inhibition in cultured hypothalamic neurons impairs FAOx (as indicated by increased palmitoyl-carnitine) and reduces oxidation of FA derived from ATGL-mediated LD lipolysis^25^. This thereby suggests that the increased activity of POMC neurons in POMC^ATGL^KO mice may involve reduced FAOx. This is supported by findings showing that mild disruption of OXPHOS increases FAOx and reduces POMC neuron activity^13^. Together, these data suggest that endogenously produced FA and FAOx inhibit POMC tone. This model is consistent with the inhibition of POMC neurons induced by exogenous palmitate or oleate during electrophysiological recordings^4^. In contrast to POMC neurons, exogenous FA^4^, impaired FAOx ^11,12^ or blockage of ATGL-dependent LD lipolysis in AgRP/NPY neurons^25^ leads to reduced neuronal activity. These findings thereby highlight a differential regulation of neuronal activity by ATGL in POMC vs. AgRP neurons.

Our data reveal that loss of ATGL in POMC neurons in males, while increasing POMC activity, does not fully reproduce the anorexigenic and catabolic effects usually induced by POMC activation^29–31^. Given that the POMC-Cre and POMC-GFP strains are known to label, in adulthood, neurons that do not actively express POMC due to transient POMC expression during development, the phenotypes observed in POMC^ATGL^KO mice may reflect alterations in LD lipolysis within a broader POMC-lineage neuronal population including AgRP/NPY neurons^32^. However, since loss of ATGL in AgRP neurons induces a distinct phenotype (i.e. no change in RQ and impaired feeding responses to cold) compared to POMC^ATGL^KO, this suggests the contribution of ATGL loss in POMC-lineage neuronal populations on mouse phenotype is minimal. Our data are in line with the study by Leon-Mercado *et al*. who found that loss of FA synthase similarly increased POMC neuron activity which was associated with decreased food intake and adiposity^14^.

Moreover, we found that loss of ATGL in POMC neurons reduces the amount of calories consumed after a fast (Fig. 3i-k), a phenotype in agreement with the anorectic effect of POMC neuron^32^. It is highly likely that reduced feeding and meal size in male POMC^ATGL^KO mice is a consequence of increased POMC firing (Fig. 1i-k)^29–31^. We also show that POMC^ATGL^KO male mice have reduced glycaemia and plasma insulin in fasted conditions, along with reduced glucose-stimulated insulin secretion during a tolerance test (Fig. 2f-k), suggesting increased insulin sensitivity, which is consistent with increased POMC activity^33–37^. Furthermore, POMC^ATGL^KO males have increased RQ in fed and refed conditions (Fig. 3c,g) demonstrating enhanced carbohydrate utilization. These results are in line with the study of Dodd *et al*. showing that chronic chemogenetic POMC activation reduces hepatic glucose production by enhancing insulin sensitivity^33,34^. Finally, in basal conditions, POMC^ATGL^KO females do not show any differences compared to POMC^ATGL^CRE mice (Fig. S2a-p), suggesting that LD lipolysis plays a different role in males *versus* females. However, these findings do not exclude an important role for LD lipolysis in females, as compensatory mechanisms or sex-specific pathways may mask the effects of ATGL loss.

Surprisingly, during fasting and cold exposure, POMC^ATGL^KO male and female mice did not show any differences in adaptive metabolic responses compared to POMC^ATGL^CRE littermates (Fig. S3a-p). One possibility is that other enzymes responsible for LD breakdown may compensate for the loss of ATGL (such as hormone sensitive lipase^38^ or DDHD2^20,21^) during metabolic challenges. Another possibility is that ATGL regulates the activity of sub-populations of POMC neurons important for glucose homeostasis but not POMC neurons that regulate EE or systemic lipolysis during a fast and cold exposure^39^. POMC neurons are involved in the regulation of thermogenesis and brown adipose tissue activation during cold exposure^39–41^. Therefore, the absence of a phenotype in POMC^ATGL^KO mice during cold exposure indicates that ATGL-dependent LD lipolysis is not required for metabolic responses to cold. Notwithstanding negative energy states, we found no beneficial effect on diet-induced obesity and glucose intolerance in POMC^ATGL^KO mice under HFD. Timper *et al*. showed that increased FAOx in POMC neurons protects against diet-induced obesity^13^. Here, our data instead suggest that blockage of LD lipolysis in POMC neurons does not play a significant role in body weight gain under HFD conditions. One possible explanation is that HFD feeding itself suppresses ATGL expression as we observed in the ARC of HFD-fed mice (Fig. S4a-b). This suggests that HFD-induced downregulation of ATGL in control mice may mask the effect of ATGL loss in KO animals. Collectively, our findings show that LD lipolysis in POMC neurons plays a significant role in neuronal activity and peripheral metabolism with strong male-biased effects. This study thereby strengthens the emerging concept that neuronal LD supports neuronal function^22,24^ including in neuronal populations that are essential for energy homeostasis^25^.

## METHODS

### Animal ethics

All animal care and experimental procedures were conducted in accordance with guidelines of the Canadian Council on Animal Care (protocol number from CIPA of CRCHUM #CM19018TAs and CM23041TAs). Mice were housed on a 12 h dark-light cycle (lights off at 10 am) at 22-23 °C, in a pathogen free environment. Standard irradiated chow diet (Teklad) and water were provided *ad libitum* or otherwise mentioned. For all studies, age- and sex-matched littermates were included and individually housed during diet interventions or phenotyping in metabolic cages.

### Primary hypothalamic cultures

The mediobasal hypothalamus from POMC-GFP P1-P2 pups (derived from POMC-eGFP reporter mice [C57BL/6J-Tg(Pomc-EGFP)1Low/J, 009593] bred with C57BL/6J WT females) was dissected in 10 mL of Lebovitz-15 media (Gibco, #11415-064) and centrifuged at 2000 g for 2 min as we described^25^. The pellet was then digested at 37 °C for 15 min using 4.5 U/mL lyophilized papain (Worthington Biochemical, #LS003120), 5 mg/mL D-glucose (Sigma, #G7528-1KG), 0.2 mg/mL L-cystein (MP, #101444) 0.2 mg/mL BSA (Multicell, #800-095-EG) and 20 U/mL DNAse I (Worthington Biochemical, #LS006333) in PBS 1X and centrifuged at 2000 g for 2 min. The pellet was then resuspended in Neurobasal A media 1X (Gibco, #10888-022) with 2 % B-27 supplement (Gibco, #17504-044), 2 % FBS (Wisent™, #10437-028), 1 % P/S (Gibco™, #15070-063) and 1 % GlutaMax (Fisher, #35050061) and filtered using a 70 μM cell-strainer. Neurons were then allowed to grow on coverslips for 7 days in Neurobasal A media with 2 % B-27, 2 % FBS, 1 % P/S and 1 % GlutaMax and 2 μM Cytosine beta-D-arabinofuranoside (AraC; Sigma, #C1768-100MG) to inhibit glial cell growth. Neurons were treated with 50 μM Atglistatin (Cedarlane, #HY-15859) or 0.1 % DMSO for 24 h, fixed using 4 % paraformaldehyde, stained using 1:1000 LipidSpot^610^ (Biotium, #70069T) for lipid droplets and 1:1500 Hoechst 33342 for nuclei (Invitrogen, #H3570) and imaged using the confocal microscopy (Carl Zeiss AG).

### Mouse studies

POMC-Cre were previously generated^42^ and males were obtained homozygous from the Jackson laboratory [Tg(Pomc1-cre)16Lowl, 005965] (FVB/N background) and bred with C57BL/6J WT females. Male POMC-eGFP reporter mice [C57BL/6J-Tg(Pomc-EGFP)1Low/J, 009593] were purchased from The Jackson Laboratory and bred with C57BL/6J WT females. ATGL^fl/fl^ mice in which exon 1 is flanked by loxP sequences were kindly donated by Dr Grant Mitchell^43^ and maintained at least 6 generations on the C57BL/6J genetic background (C57BL/6J, Jackson Lab #000664).

#### ATGL KO in POMC neurons (POMC^ATGL^CRE vs POMC^ATGL^KO)

The Cre-Lox system was used to excise exon 1 of the ATGL floxed gene in POMC neurons. The Cre locus was maintained hemizygous, to avoid potential Cre toxicity. Briefly, mice POMC^Cre/+^ were bred with POMC-GFP. Male POMC^Cre/+^:POMC^GFP/+^ were bred with female ATGL^fl/fl^ mice to obtain POMC^Cre/+^:POMC^GFP/+:^ATGL^fl/+^ mice which were then crossed with ATGL^fl/+^ or ATGL^fl/fl^ to generate experimental mice: POMC^Cre/+^:ATGL^+/+^ (POMC^ATGL^CRE) and POMC^Cre/+^:ATGL^fl/fl^ (POMC^ATGL^KO). These experimental mice were used independently of the genotype for POMC^GFP/+^ or POMC^+/+^. Mice harboring significant ectopic ATGL recombination in ear punch tissue (detected by PCR on gel) were excluded from experimentations (N = 6 mice with significant ectopic recombination).

#### High fat diet studies

Animals were exposed to a palm-enriched high fat-diet (HFD) in which 50 % kcal are derived from palm oil (Dyet, Bethlehem, PA, USA). Briefly, male C57BL/6J were fed ad *libitum* with a HFD for 12 weeks starting at 56 days of age. POMC^ATGL^CRE and POMC^ATGL^KO males were fed ad *libitum* with the HFD diet over the course of 8 weeks starting at 56 days of age. POMC^ATGL^CRE and POMC^ATGL^KO females were subjected to *ad libitum* HFD diet over the course of 12 weeks starting at 56 days of age.

#### Metabolic cages (CLAMS)

RQ, EE, food consumption, FAOx and locomotor activity were monitored using indirect calorimetry in Comprehensive Lab Animal Monitoring System metabolic cages (CLAMS, Columbus Instruments International). Animals were single-housed in CLAMS cages in a dark/light cycle matching their housing conditions during 24 h for acclimation, followed by measurements. For fasting-refeeding experiments: mice were food deprived for 16 h starting during the second half of the dark period and a full light period to then regain access to food *ad libitum* during 24 h. For cold exposure experiments, body temperature was monitored with implantable transmitters that were surgically inserted intra abdominally and mice recovered for at least 2 weeks. Measurements included 24 h at 21 °C followed by 24 h at 4 °C.

#### Feeding behavior analysis

The CLAMS system weighs hoppers with food (± 0.01 g) every second and detects “not eating” when weight is stable and “eating” if unstable. Single feeding events (bouts) are calculated as the weight difference between “eating” and “not eating” events as we described^25^. Data for single mice were extracted using Oxymax software then analyzed and compiled with MATLAB (MathsWorks© R2021a). As previously reported, meals consist of the sum of single feeding events (bouts) separated by an inter-meal interval (IMI)^44,45^. A meal is the sum of bouts >0.03 g and >10 sec, occurring within <5 min. For a defined period of time, following parameters were calculated using the MATLAB program: meal number, cumulative food intake (the sum of all meals (g)), and average meal size (g).

#### Body composition

Body composition (fat and lean mass) was measured by magnetic resonance imaging (echoMRI). Brown adipose tissue (BAT), inguinal, intraperitoneal (perigonadal) and subcutaneous (inguinal) fat pads were collected and weighed using an analytical scale (Sartorius) at sacrifice.

#### Glucose tolerance test

Mice were food deprived 5 h before the test at ZT10. A bolus of glucose (Dextrose, 1.5 g/kg) was administered via an intraperitoneal injection, and glycemia was measured from blood sampled from the tail vein using an Accu-chek Performa glucometer at T0 (before injection), 15, 30, 60, and 90 min. Tail vein blood samples were collected via a capillary for insulin assays.

#### Insulin assay

Insulin assays were performed by the metabolic phenotyping core facility of the CRCHUM using commercially available ELISA kits.

#### Stress restraint and corticosterone assay

Tail blood was collected between ZT11-12 in capillary tubes and centrifuged at 8000 rpm for 5 min to retrieve plasma. Plasma corticosterone was measured at time zero and 15 min following a restraint stress using Enzo Corticosterone ELISA kits (#ADI-900-097, Enzo).

#### Tissue collection

Fresh brain microdissection (ARC) and tissue collection were performed in deeply anesthetized mice (isoflurane). Tissues were weighed, flash frozen and stored at -80 °C.

Perfusions were performed for cytomorphologic experiments. Mice were deeply anesthetized with excess Ketamin/xylazin and transcardially perfused with 1x PBS followed by 4 % PFA. Brains were extracted, post-fixed for 2 h in 4 % PFA, placed in sucrose overnight and stored at -80 °C. Using a microtome (Leica, SM2000R), 30*μ*m brain coronal sections were made and kept in antifreeze at -20 °C before any imaging.

#### LD detection in brain sections

Brain sections (30 µm) were washed twice for 5 min in 1X PBS and stained with 1:1000 LipidSpot^610^ (VWR, 70069-T) in 1X PBS, and 1:5000 Hoechst 33342, 2 h at room temperature. Sections were washed twice in 1X PBS for 5 min and mounted using ProLong™ Gold Antifade Mountant (eLife, #P36930). Imaging was performed using a Leica Stellaris 8 confocal microscope on a 63x oil-immersion lens, with 0.3 µm z-steps (with an average of 70 z-steps per neuron) to create 3D reconstruction of soma neurons and LD with Imaris software as we previously described^25^. On Imaris, LD were defined as: Spots with an average diameter of 0.4 µm as per indicated in previous *in vitro* studies on LD in brain cells^46^, background substracted, z-diameter for model PSF elongation of 1.2, quality of 260, and region threshold for local contrast of 32.4.

#### Gene expression

Total RNA from frozen tissue was extracted using the Trizol method as previously described^47^. Total RNA concentration and purity were determined using the Nanodrop 2000. cDNA synthesis and qPCR were performed as described^48^. Briefly, 1 µg of total RNA was retro-transcribed with M-MuLV reverse transcriptase (Invitrogen, #28025013) using random hexamers and diluted 1:10 prior to quantification by qPCR (QuantiFast SYBR Green PCR kit, Qiagen, #28025013) using a Corbett Rotor-Gene 6000 (Qiagen) with primers (1 µM) as described here (forward, reverse) : 18S (TAGCCAGGTTCTGGCCAACGG, AAGGCCCCAAAAGTGGCGCA), B-actin (TTCTTGGGTATGGAATCCTGTGGCA, ACCAGACAGCACTGTGTTGGCATA), Cyclophilin (GCTTTTCGCCGCTTGCTGCA, TGCAAACAGCTCGAAGGAGACGC), mATGL (TCACCATCCGCTTGTTGGAG, GAAGGCAGATGGTCACCCAA), POMC(CAGTGCCAGGACCTCACC, CAGCGAGAGGTCGAGTTTG). qPCR was quantified using the standard curve method. Gene expressions were normalized to the expression of a stable housekeeping gene or the geometric mean of many (determined by NormFinder^49^). Gene expressions are represented as fold change from mean of control groups, after normalization.

#### RNAScope

Brain sections (30 *μ*m) were washed with 1X PBS, mounted on microscope glass slides, and dried for 30 min at 60 °C. Following the company’s instructions (ACD, Multiplex Fluorescent V2 Assay, #323100), on the 1st day, all sections were dehydrated, incubated with hydrogen peroxide, steamed for 5 min with antigen retrieval, and treated with Protease III for 15 min. Pnpla2 (#469441-C1) and/or POMC (#314081-C2) probes were incubated 2 h at 40 °C as indicated (C2 probe diluted at 1:50 in the c1-probe), and a negative control brain section, was done for every sample using a negative probe (ACD, #320871). Slides were kept ON, in 5x Saline Sodium Citrate protected from light, at RT. On the second day, after washing in Wash Buffer, sections were amplified 3 times as indicated by the company. Fluorescent signal was developed using opal dye diluted at 1:1500, sections were counterstained with Dapi and mounted with ProLong Gold Antifade Mountant (eLife, #P36930) and stored at 4 °C. Zeiss fluorescent microscope (Carl Zeiss AG) with an Apototome was used for imaging. Quantification was done using Fiji software.

#### Electrophysiology

Seven weeks old POMC^ATGL^CRE and KO mice underwent stereotaxic surgery to selectively label POMC neurons with the CRE-dependent red mCherry fluorescent protein as we described^25,50^. Briefly, mice were kept under anesthesia with isoflurane and received bilateral viral injections of AAV2-CAG-DIO-mCherry-WPRE-bGHpA (Viroveck, 1.31E13 vg/ml) in the ARC. 200 nL/side were simultaneously injected (0.5 nl/sec) using neurosyringes (Hamilton, #65457-01) placed at a 10° angle, to the following coordinates AP: bregma-1.4mm ; lateral: sinus+1.2mm ; depth: dura-5.8 mm. Syringes were removed 5 min after the injection, the wound closed and mice recovered in thermoneutrality incubators for at least 2 h.

##### ARC slices

Coronal slices (300 µm thick) were obtained from POMC^ATGL^CRE and POMC^ATGL^KO mice 3 to 4 weeks after the surgeries. Animals were deeply anesthetized with isoflurane and killed by decerebration. Then the brain was rapidly removed and cut with a vibratome VT1200 (Leica, Germany) in ice-cold N-methyl-D-glucamine (NMDG) cutting ACSF containing (in mM): 119.9 NMDG, 2.5 KCl, 25, 1 CaCl_2_, 1.4 NaH_2_P0_4_ and 20 D-glucose saturated with 95 % O_2_ and 5 % CO_2_. Slices containing the ARC were transferred to 95 % O_2_ and 5 % CO_2_ saturated ASCF containing (in mM): 2.5 glucose, 130 NaCl, 2.8 KCl, 1.25 NaH_2_PO_4_, 1.2 MgCl_2_, 2.5 CaCl_2_ for 30 min at 32°C before a recovery period of at least 20 min at room temperature prior to recordings.

##### Recording

Slices were transferred to a recording chamber where they were perfused with ACSF (2 ml/min) containing 2.5 mM glucose. Whole-cell recordings were achieved from the soma of mCherry expressing ARC neurons. Borosilicate patch pipettes (2-5 MΩ) were filled with an internal solution containing (in mM): 105 K-gluconate, 30 KCl, 10 phosphocreatine, 10 HEPES, 4 ATP-Mg, 0.3 GTP-Tris, and 0.3 EGTA (adjusted to pH 7.2 with KOH; 290-300 mOsmol). Recordings were made at room temperature and only a single neuron was recorded per slice. Data were acquired using a Multiclamp 700B amplifier (Molecular Devices) and digitized using a Digidata 1440A digitizer and pClamp/Clampfit 10.7 (Molecular Devices). Recordings were low pass-filtered at 2 kHz and digitized at 20 kHz. Access resistance (Ra) was 10-25 MΩ and regularly monitored during experiments, data were excluded if Ra variations were above 20 % throughout the experiment. Spontaneous firing frequencies were obtained from I=0 current-clamp mode recordings for at least 2 min.

## ACKNOWLEDGMENTS

We acknowledge Dr Grant Mitchell (U. Montréal) for the ATGL-floxed mice. We thank Dr Xavier Fioramonti (U. Bordeaux) for his help in optimizing hypothalamic electrophysiology studies. We acknowledge the cellular imaging, small animal phenotyping, cell physiology and imaging core facilities at the CRCHUM. This work was supported by grants to TA from the Natural Sciences and Engineering Research Council of Canada (NSERC, RGPIN-2025-04602) and the Canadian Institutes of Health Research (CIHR, PJT178131). SF, CMR and TA were supported by a salary award from Fonds de Recherche Québec-Santé (FRQS). JLR was supported by a postdoctoral fellowship from FRQS. DM, RM, LRDH and FB were supported by doctoral fellowships from FRQS, Diabète Québec, the Neuroscience Department and Faculty of Graduate and Postdoctoral Studies of Université de Montréal, and CRCHUM.

## AUTHOR CONTRIBUTIONS

DM and MFF performed studies on cultured mouse neurons. DM, RM, FB, DR, JR, KB conducted mouse colony management, genotyping, RNAscope, histology, qPCR, stereotaxic injections and phenotyping studies. LRDH and CMR carried out electrophysiological recordings and data analysis. DM, RM, FB, MFF, JR, SF, CMR and TA contributed to conceptualization, experimental design for cultured neuron and mouse studies, data analysis and interpretation. DM, FB, RM and TA drafted and edited the manuscript.

