## Supplemental Figures 1 to 4 for "Lipid droplet lipolysis in POMC neurons regulates energy homeostasis in a sex-specific manner"

### Figure S1

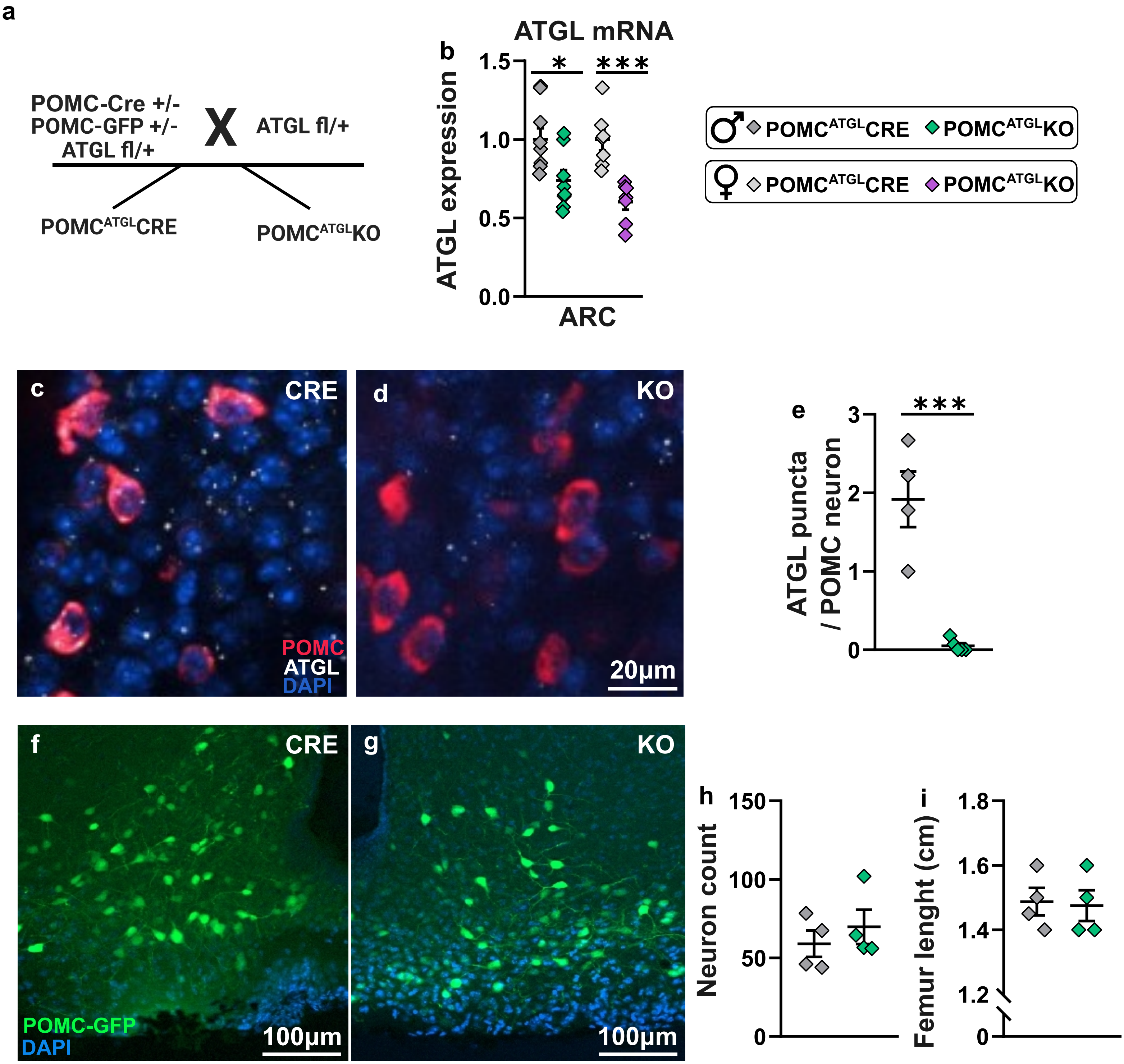

a
POMC-Cre +/-
POMC-GFP +/-
ATGL fl/+
X
ATGL fl/+
POMCATGLCRE
POMCATGLKO
b
ATGL mRNA
1.5
1.0
0.5
0.0
ATGL expression
*
***
ARC
♂ POMCATGLCRE POMCATGLKO
♀ POMCATGLCRE POMCATGLKO
c
CRE
POMC
ATGL
DAPI
d
KO
20µm
e
3
2
1
0
***
ATGL puncta / POMC neuron
f
CRE
POMC-GFP
DAPI
100µm
g
KO
100µm
h
150
100
50
0
Neuron count
i
1.8
1.6
1.4
1.2
0
Femur lenght (cm)

**Figure S1. POMC$^{ATGL}$KO model validation.** Validation of the POMC-Cre x ATGL LoxP mouse model (**a**) using qPCR (**b**) and RNAscope (**c-e**). N = 4-7 mice/genotype. POMC-GFP neuron number (**f-h**) and femur length in POMC$^{ATGL}$KO and POMC$^{ATGL}$CRE male mice (**i**). N = 4 mice/genotype. Data are represented as mean ± SEM. Student's t-test. *$p<0.05$, ***$p<0.001$.

### Figure S2

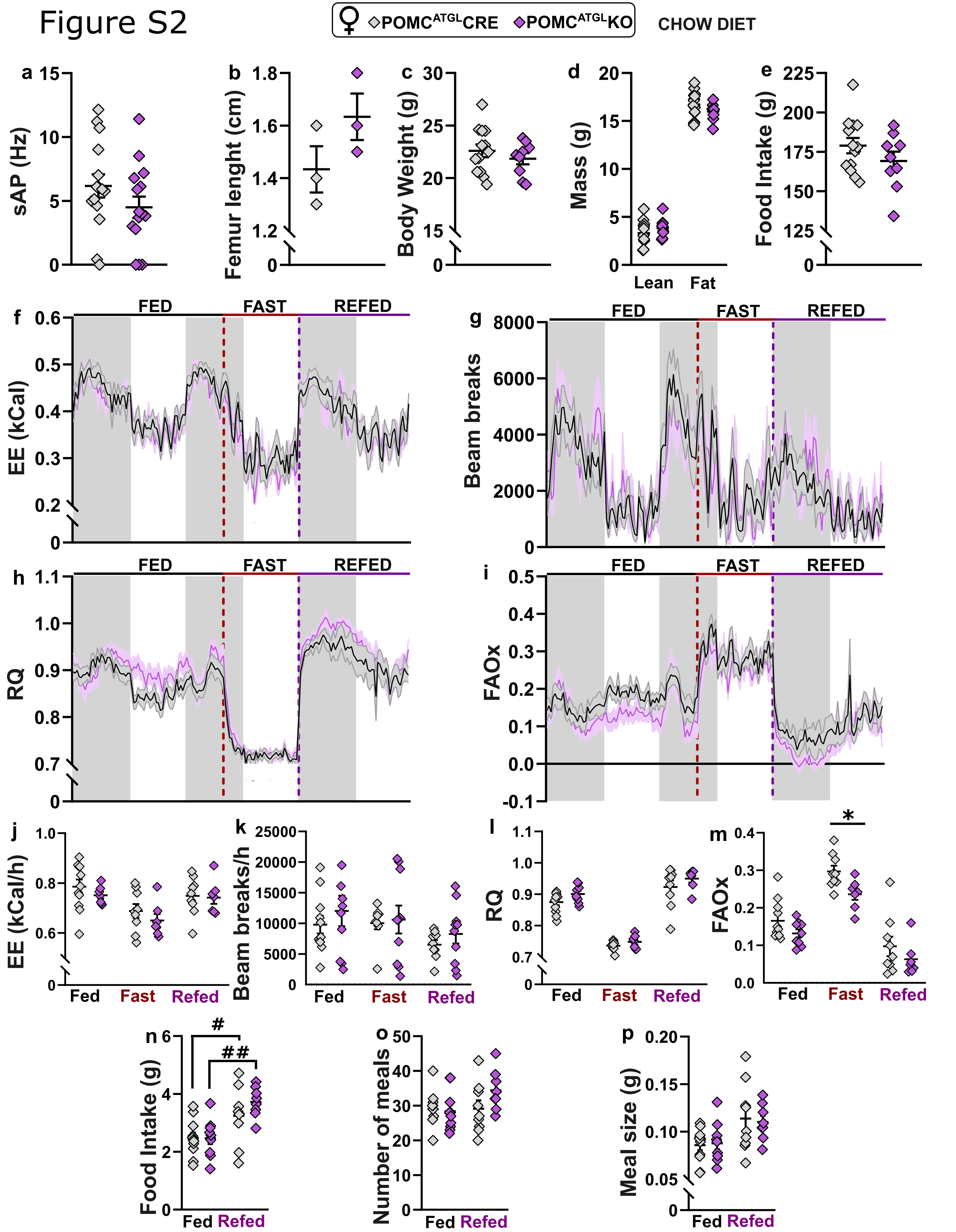

♀ POMCATGLCRE POMCATGLKO
CHOW DIET
a
sAP (Hz)
b
Femur lenght (cm)
c
Body Weight (g)
d
Mass (g)
Lean
Fat
e
Food Intake (g)
f
EE (kCal)
FED
FAST
REFED
g
Beam breaks
h
RQ
i
FAOx
j
EE (kCal/h)
Fed
Fast
Refed
k
Beam breaks/h
l
m
*
n
Food Intake (g)
#
##
o
Number of meals
p
Meal size (g)

**Figure S2. POMC$^{ATGL}$KO females do not show changes in POMC neuron activity nor energy balance under chow diet.** Frequency of spontaneous action potential of POMC neurons (**a**), body weight (**b**), body composition (**c**), food intake (**d**), and femur length (**e**) of POMC$^{ATGL}$KO and POMC$^{ATGL}$CRE females under 8 weeks of standard chow diet. N = 3-13 mice/genotype. Parameters of energy balance measured in metabolic cages including EE (**f, j**), locomotor activity (**g, k**), RQ (**h, l**), and FAOx (**i, m**) in POMC$^{ATGL}$KO and POMC$^{ATGL}$CRE females during 24 h in chow-fed conditions, a 16 h fast and 24 h refeeding. N = 5-8 mice/genotype. Food intake (**n**), number of meals (**o**), and meal size (**p**) in POMC$^{ATGL}$KO and POMC$^{ATGL}$CRE females during fed and refed conditions. N = 5-8 mice/genotype. RQ; Respiratory quotient, FAOx; Fatty acid oxidation, EE; Energy expenditure. Data are represented as mean ± SEM. Student's t-test (**a-e, n-p**) or two-way ANOVA with Sidak's post-hoc test (**f-m**); * indicates a genotype effect while # indicates time effect. *$p<0.05$, $^{\#}p<0.05$, $^{\#\#}p<0.01$.

### Figure S3

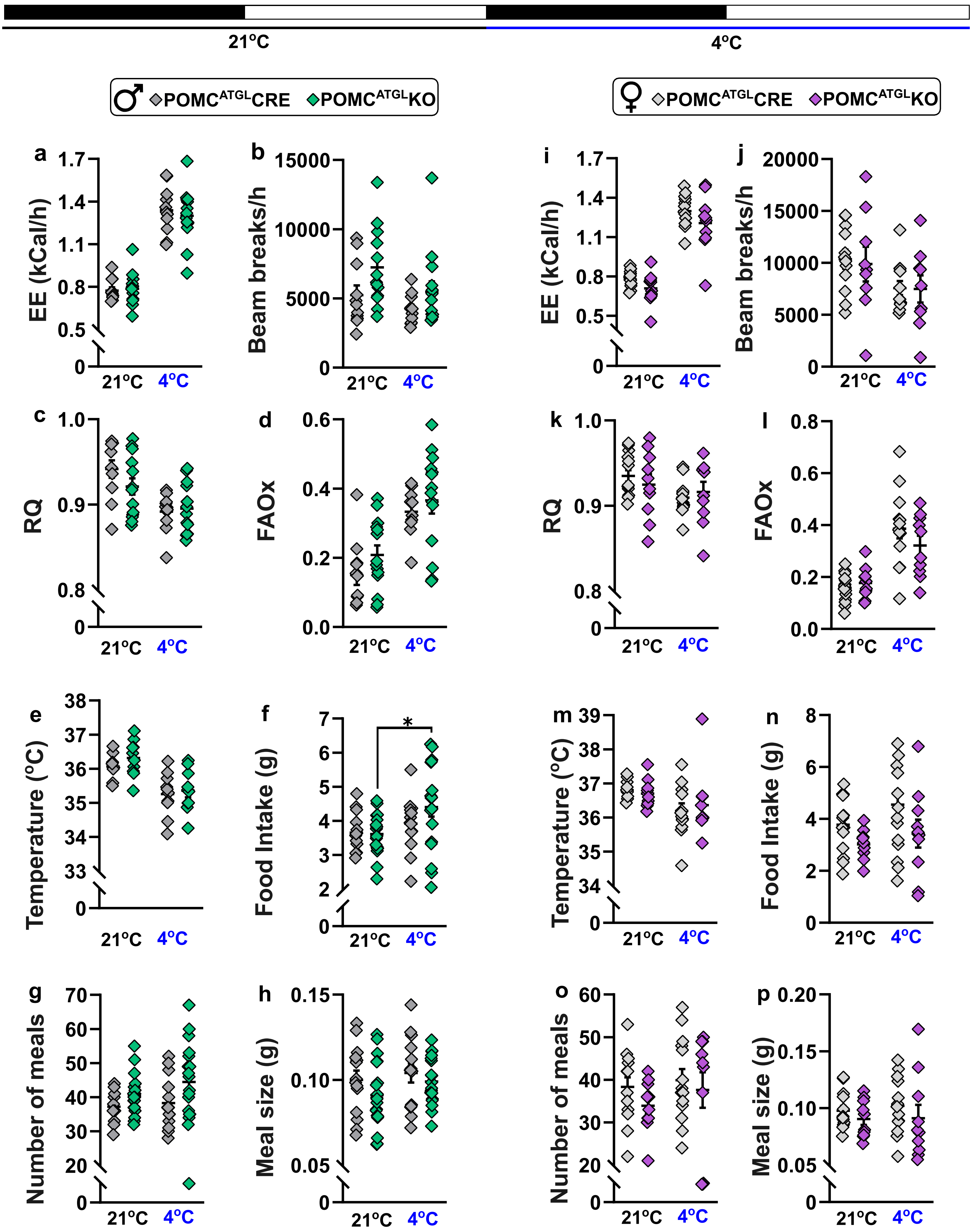

21°C
4°C
POMC^ATGL CRE
POMC^ATGL KO
a
EE (kCal/h)
b
Beam breaks/h
c
RQ
d
FAOx
e
Temperature (°C)
f
Food Intake (g)
*
g
Number of meals
h
Meal size (g)
i
j
k
l
m
n
o
p

**Figure S3. Adaptive responses to cold are not affected in POMC$^{ATGL}$KO males and females.** Measures of metabolism including EE (**a**), locomotor activity (**b**), RQ (**c**), FAOx (**d**), core body temperature (**e**), food intake (**f**), number of meals (**g**), and meal size (**h**) in male POMC$^{ATGL}$KO and POMC$^{ATGL}$CRE during 24 h at 21ºC and 24 h cold exposure at 4ºC. N = 10-14 mice/genotype. Measures of metabolism including EE (**i**), locomotor activity (**j**), RQ (**k**), FAOx (**l**), core body temperature (**m**), food intake (**n**), number of meals (**o**), and meal size (**p**) in female POMC$^{ATGL}$KO and POMC$^{ATGL}$CRE during 24 h at 21ºC and 24 h cold exposure at 4ºC. N = 10-13 mice/genotype. Data are represented as mean ± SEM. Two-way ANOVA with Sidak's post-hoc. * represents a genotype effect. **p<0.01.

### Figure S4

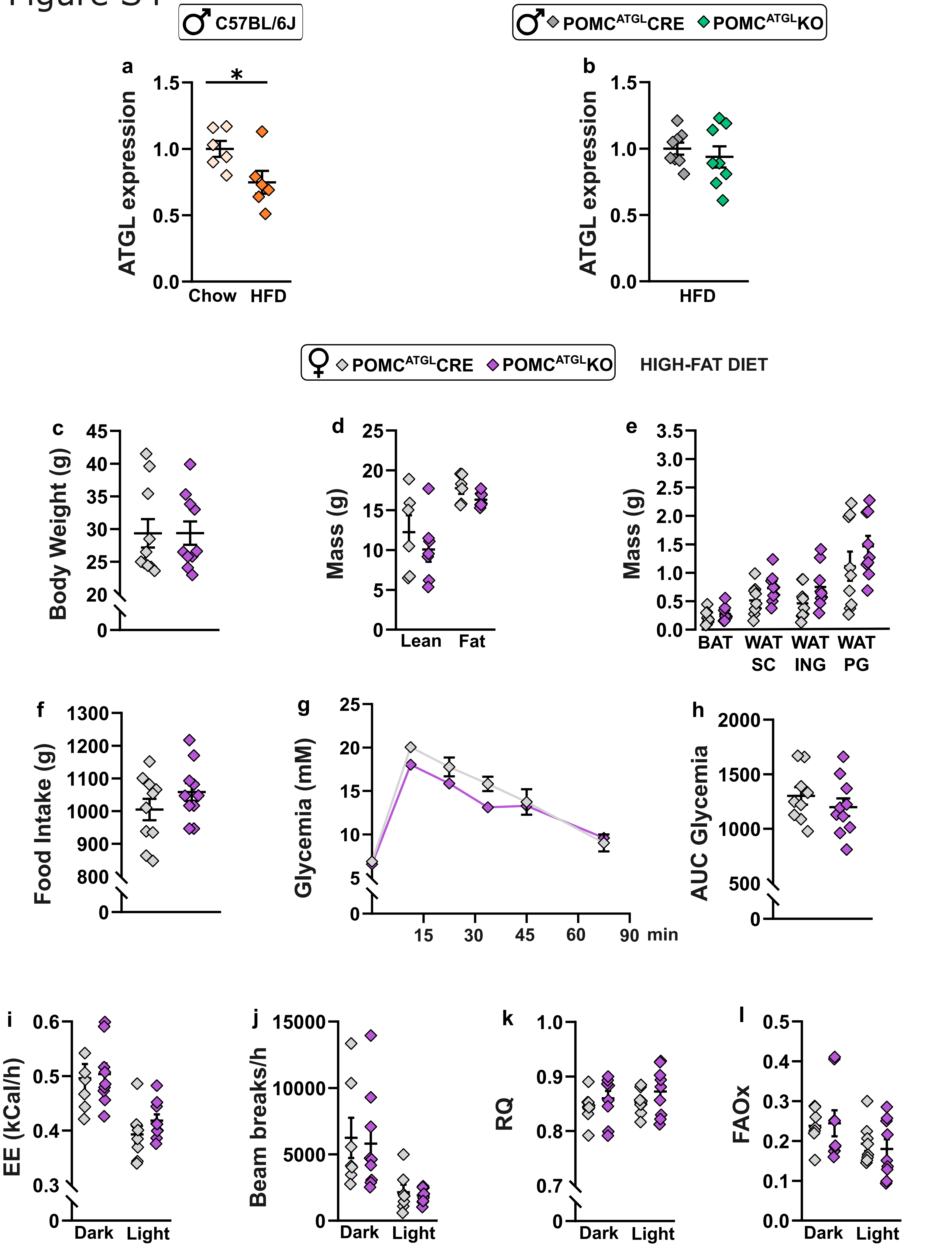

C57BL/6J
POMC^ATGL CRE
POMC^ATGL KO
a
b
ATGL expression
1.5
1.0
0.5
0.0
*
Chow
HFD
HFD
POMC^ATGL CRE
POMC^ATGL KO
HIGH-FAT DIET
c
Body Weight (g)
45
40
35
30
25
20
0
d
Mass (g)
25
20
15
10
5
0
Lean
Fat
e
Mass (g)
3.5
3.0
2.5
2.0
1.5
1.0
0.5
0.0
BAT
WAT SC
WAT ING
WAT PG
f
Food Intake (g)
1300
1200
1100
1000
900
800
0
g
Glycemia (mM)
25
20
15
10
5
0
15
30
45
60
90
min
h
AUC Glycemia
2000
1500
1000
500
0
i
EE (kCal/h)
0.6
0.5
0.4
0.3
0
Dark
Light
j
Beam breaks/h
15000
10000
5000
0
Dark
Light
k
RQ
1.0
0.9
0.8
0.7
0
Dark
Light
l
FAOx
0.5
0.4
0.3
0.2
0.1
0.0
Dark
Light

**Figure S4. ATGL expression in the ARC of HFD-fed males and loss of ATGL in POMC neurons does not affect diet-induced obesity in females.** (**a**) ATGL expression in the ARC of WT males fed during 12 weeks with a high-fat diet. N = 6 mice/condition. (**b**) ATGL expression in the ARC of $POMC^{ATGL}KO$ and $POMC^{ATGL}CRE$ males fed with a high fat diet during 12 weeks. N = 8 mice/genotype. Body weight (**c**), body composition (**d**), fat pad weight (**e**), food intake (**f**), and glycemia during intra-peritoneal glucose tolerance test (**g-h**) in $POMC^{ATGL}KO$ and $POMC^{ATGL}CRE$ females after 12 weeks of high-fat diet. N = 6-7 mice/genotype. Parameters of energy balance measured in metabolic cages including EE (**i**), locomotor activity (**j**), RQ (**k**), and FAOx (**l**) in high-fat diet fed $POMC^{ATGL}KO$ and $POMC^{ATGL}CRE$ females during 24 h. N = 8-9 mice/genotype. RQ; Respiratory quotient, FAOx; Fatty acid oxidation, EE; Energy expenditure, AUC; Area under the curve. Data are represented as mean ± SEM. Student's t-test (**a-f, h**) or two-way ANOVA with Sidak's post-hoc test (**g, i-l**). *$p<0.05$.
